# Platypus Induced Pluripotent Stem Cells: the Unique Pluripotency Signature of a Monotreme

**DOI:** 10.1101/377010

**Authors:** DJ Whitworth, IJ Limnios, M-E Gauthier, P Weeratunga, DA Ovchinnikov, G Baillie, SM Grimmond, JA Marshall Graves, EJ Wolvetang

## Abstract

**Summary Statement:** The generation and transcriptome analysis of the first induced pluripotent stem cells from the platypus reveals SOX2 has been a key driver of the expanded pluripotency regulatory network in placental mammals.

**ABSTRACT:** The mechanisms by which pluripotency has evolved remain unclear. To gain insight into the evolution of mammalian pluripotency we have generated induced pluripotent stem cells (piPSCs) from the platypus. Deep sequencing of the piPSC transcriptome revealed that piPSCs robustly express the core eutherian pluripotency factors *OCT4, SOX2* and *NANOG*. Given the more extensive role of *SOX3* over *SOX2* in avian pluripotency, our data indicate that between 315 million years and 166 million years ago primitive mammals replaced the role of *SOX3* in the vertebrate pluripotency network with *SOX2. DAX1/NR0B1* is not expressed in piPSCs and an analysis of the platypus *DAX1* promoter revealed the absence of a proximal SOX2-binding DNA motif known to be critical for *DAX1* expression in eutherian pluripotent stem cells, suggesting that the acquisition of SOX2 responsiveness by *DAX1* has facilitated its recruitment into the pluripotency network of eutherians. We further show that the expression ratio of X chromosomes to autosomes (X_1-5_ X_1-5_:AA) is approximately equal to 1 indicating that there is no upregulation of X-linked genes and that there is no preference for silencing of maternal or paternal alleles (ie imprinting).

## INTRODUCTION

The platypus *(Ornithorhynchus anatinus)* is an extraordinary mammal: in addition to its unique, and defining, toothless bill, females lay reptilian-like eggs while nourishing their young through lactation; they use electroreception to detect aquatic prey, and males produce a complex venom (De Plater et al. 1995). This amalgamation of ancestral reptilian and derived mammalian traits is similarly reflected in its genome (Warren et al. 2008). The platypus belongs to the Order Monotremata, that also includes several species of echidna, which is thought to have diverged from the therian lineage that gave rise to marsupial (Infraclass Marsupialia) and eutherian (Infraclass Placentalia) mammals approximately 166 million years ago (Bininda-Emonds et al. 2007). Thus, the platypus affords the opportunity to examine the emergence and evolution of mammalian traits. In particular, we wished to use the platypus to better understand the evolution of pluripotency in mammals.

The acquisition and maintenance of pluripotency by a small subpopulation of cells within the mammalian embryo requires a complex series of tightly regulated interactions between the factors that comprise the pluripotency network, or plurinet. In eutherian mammals, the core pluripotency factors *OCT4, SOX2* and *NANOG* together regulate other downstream members of the plurinet (Boyer et al. 2005; Loh et al. 2006). To date, very little is known about the evolutionary origins of the plurinet and its conservation amongst the vertebrates. In the chicken, *POU2/POUV* (a paralogue of mammalian *OCT4)* and *NANOG* similarly constitute the core components of the avian plurinet (Lavial et al., 2007; Fernandez-Tresguerres et al. 2010; Jean et al., 2015); the role of *SOX2,* however, is less clear.

Monotremes are very strategically placed within this 315 million year divergence, sitting at the cusp of mammalian evolution. With their combination of ancestral reptilian-like traits and derived mammalian characteristics, an examination of the monotreme plurinet should enable us to identify some of the genetic events that have contributed to the assembly of the mammalian pluripotency network. Our data indicate that the platypus shares a greater proportion of pluripotency factors with eutherian mammals than with the chicken, but has many more pluripotency-related genes in common with the chicken than do any of the eutherian species examined. In combination with our recent data on pluripotency in a marsupial (Weeratunga et al., 2018), this study supports the argument that the evolution of pluripotency in mammals has involved expanding the role of *SOX2* over *SOX3,* and as regards specifically eutherian pluripotency, *DAX1/NR0B1* and *ESRRB* have been recruited into the plurinet likely through the acquisition of SOX2 responsiveness.

## MATERIALS AND METHODS

All use of animals, and tissues obtained from animals, was approved by both the Australian National University and the University of Queensland Animal Ethics Committees.

### Generation and Maintenance of Platypus iPSCs

Approximately 5 x 10^6^ fibroblasts from an adult female platypus were transduced with lentiviruses expressing human *OCT4, SOX2, KLF4, cMYC, NANOG* and *LIN28* (Addgene plasmids 21162 and 21164) as previously described (Whitworth et al. 2012). Forty-eight hours after transduction, fibroblasts were passaged onto 10 cm culture plates (Costar) coated with Matrigel (BD Biosciences) at a density of 5.0 x 10^5^ cells per plate and maintained in medium consisting of KnockOut DMEM (Gibco), 10% (v/v) KnockOut serum replacement (KOSR) (Gibco), 0.1 mM non-essential amino acids (NEAA) (Gibco), 2 mM L-glutamine (Gibco) and 0.1 mM β-mercaptoethanol (Gibco) supplemented with 1,000 U/ml murine LIF (ESGRO, Millipore), 10 ng/ml human bFGF (Invitrogen), 3 *μ*M GSK3β inhibitor (CHIR99021, Stemgent), 0.5 *μ*M MEK inhibitor (PD0325901, Stemgent), 0.25 *μ*M TGF-β antagonist (A83-01, Stemgent) and 2.5 *μ*M ALK receptor inhibitor (SB431542, Stemgent). Cultures were maintained at 32°C with 5% O_2_ and 5% CO_2_.

Putative piPSCs were subcultured onto feeder layers of irradiated mouse embryonic fibroblasts (MEFs) once the colonies were large enough to be mechanically isolated. After subculture onto MEFs, piPSCs were maintained in Knockout DMEM, 10% (v/v) KOSR, 0.1 mM NEAA, 2 mM L-glutamine and 0.1 mM β-mercaptoethanol with either 1,000 U/ml murine LIF and 10 ng/ml human bFGF, 10 ng/ml human bFGF alone or 1,000 U/ml murine LIF only. In spite of multiple reprogramming attempts, only two clones of putative piPSCs were generated, both from the same reprogramming experiment. Only one clone displayed the hallmarks of full reprogramming and was characterised further.

### Karyotyping

PiPSCs were commercially karyotyped by Sullivan Nicolaides Pathology (Taringa, QLD, Australia).

### Alkaline Phosphatase Staining and Fluorescence Immunocytochemistry

Alkaline phosphatase staining was performed using the Quantitative Alkaline Phosphatase Detection Kit (Millipore) as per the manufacturer’s instructions.

Fluorescence immunocytochemistry was performed for the following markers of pluripotency: SSEA1, SSEA4, TRA1-60 and TRA1-81. Samples were fixed with 4% paraformaldehyde at RT for 10 minutes before blocking with 5% goat serum (Gibco) in phosphate buffered saline (PBS) (Gibco) and incubated in primary antibody diluted in 3% goat serum in PBS. Primary antibodies, and their dilutions, are as follows: anti-SSEA1 (MAB4301, Millipore), 1:50; anti-SSEA4 (MAB4304, Millipore), 1:50; anti-TRA1-60 (MAB4360, Millipore), 1:50 and anti-TRA1-81 (MAB4381, Millipore), 1:50. Negative controls were incubated with 3% goat serum in PBS in place of primary antibody. All secondary antibodies were diluted to 1:1000 in PBS: Alexa Fluor goat anti-mouse IgG_H&L_ (Invitrogen) and Alexa Fluor goat anti-rabbit IgG_H&L_ (Invitrogen).

The rate of cell proliferation was determined by fluorescence immunocytochemistry using an antibody to phospho-histone H3 (Ser10) (anti-phospho-histone H3 (Ser10), 9708, Cell Signalling Technology) which identifies the phosphorylation of H3 in cells undergoing mitosis. Mitotic figures were counted in 30 high power fields (HPF) (x1000 magnification) per sample, with the samples de-identified to the experimenter, and the data represented as the mean number of mitotic figures per HPF ± standard error of the mean (SEM). Statistical significance was determined by a Student’s t test.

### RNA Isolation, cDNA Synthesis and PCR

Total RNA was isolated using the Qiagen RNeasy Mini kit (Qiagen) as per the manufacturer’s protocol. Complementary DNA was synthesised using the Bio-Rad iScript Reverse Transcriptase kit (Bio-Rad Laboratories) according to the manufacturer’s instructions. The PCR primers used, and their product size, are listed in Supplementary Table 1. Negative control reactions incorporated template from cDNA reactions that were performed without reverse transcriptase or using water in place of template in PCR reactions.

### Deep Sequencing of the piPSC Transcriptome

Reads were filtered using FastqMcf (Ea-utils) and quality-checked using FastQC. We then performed alignment and mapping on the filtered reads using the Tuxedo suite. To account for any potential contamination from MEFs, reads that aligned to the genome of *Mus musculus* (ENSEMBL build37.1) were excluded. The remaining sequences were aligned to the genome of *Ornithorhynchus anatinus* (OANA5). As the platypus genome is very fragmented (with 201,523 contigs and chromosomes), we only retained contigs predicted to encode genes by ENSEMBL (approximately 7%), as well as by NCBI for homologues of the plurinet that had no ENSEMBL prediction (eg *NANOG, POU2*-related, *SHH, IRS1, GDF9, GADD45GIP1, PIM3, PIM1, BMP4, TLE2, HCK, TERT, MAPK1, PRKCC* and *KDM4C*). Mapping was performed using TopHat (using the very sensitive parameter and setting the number of mismatches allowed in a seed alignment to 1). After running TopHat, the obtained bam files were filtered for any low quality mapped reads (MAPQ<9) as well as any matches to rRNAs and MtRNAs. We then derived FPKM values using Cufflinks (using both the fragment bias correction and the multiread correction parameters). Cufflinks was first run using the existing ENSEMBL annotation augmented by the NCBI gene prediction models of interest. We then ran Cufflinks in the discovery mode to evaluate the accuracy of the current ENSEMBL annotation and to detect novel transcripts. Genes were defined as expressed if the FPKM ≥ 1 and as differentially expressed if the FC_log_2_ ≤ -2 or ≥ 2.

### Generation of Embryoid Bodies and In Vitro Teratomas

To generate embryoid bodies (EBs), colonies were passaged with trypsin (TrypLE, Gibco) to produce small aggregates of cells which were then cultured in Costar Ultra-low Attachment plates (Corning Life Sciences) in KnockOut DMEM supplemented with 15% FCS (JRH Biosciences), 0.1 mM NEAA and 2 mM L-glutamine. After 6 weeks they were harvested for total RNA extraction and their ability to form derivatives of the 3 germ layers was assessed by RT-PCR. The PCR primers used, and their product size, are listed in Supplementary Table 1.

Because the core body temperature of the platypus is 32°C, and platypus cells do not survive at temperatures above this, the use of mice for the generation of conventional teratomas was not possible. However, we have devised and fully characterised a methodology for generating teratomas in vitro (Whitworth et al. 2012; Whitworth et al., 2014a; Whitworth et al., 2014b). While originally this protocol was applied to human ESCs and iPSCs, it has worked equally as well with horse, dog and Tasmanian devil iPSCs (Whitworth et al. 2012; Whitworth et al., 2014a; Whitworth et al. 2014b; Weeratunga et al. 2018) and so seemed applicable also for our platypus iPSCs. PiPSC colonies were passaged with trypsin (TrypLE) to produce small aggregates of cells. Approximately 4 x 10^5^ cells were sandwiched between two layers of 10% (w/v) low molecular weight methylcellulose (Sigma-Aldrich) in KnockOut DMEM. Cells were maintained in KnockOut DMEM supplemented with 20% FCS, 0.1 mM NEAA, 2 mM L-glutamine and 25 mM HEPES (Gibco) for 8 weeks before embedding and sectioning for histological analysis.

### Sequence Analysis

Genomic organisation and sequences were obtained from the Ensembl and NCBI genome browsers (www.ensembl.org and www.ncbi.nlm.nih.gov). Multiple sequence alignments were performed using ClustalW available at www.genome.jp

### Venn Analysis

Venn analysis was performed using the chicken ESC dataset from Jean et al. (2015) and the mouse and pig iPSC datasets of Liu et al. (2014), using the Venny tool at http://bioinfogp.cnb.csic.es/tools/venny/.

### Assessment of Expression Ratios of X Chromosomes to Autosomes

RNAseq data was used to calculate the X chromosome to autosome expression ratios (X_1-5_X_1-5_:AA) for both fibroblasts and iPSCs. For fibroblasts, the FPKM values for 280 X-linked genes and 11,923 autosomal genes, that were determined as being expressed (ie FPKM ≥ 1), were used to calculate median expression levels. Similarly, 299 expressed X-linked genes and 12,897 autosomal genes were used to calculate their respective median expression levels in piPSCs.

### Determination of Allelic Bias in X-Linked Transcripts From Fibroblast RNAseq Data

Previous data has shown that X-linked genes in the platypus are inactivated on a gene-by-gene basis; that is, there is no whole X chromosome inactivation in the platypus (Deakin et al., 2008; Rens et al., 2010; Livernois et al., 2013). Rather, for any given X-linked gene, a specific proportion of nuclei within a cell population will silence one locus, with the percentage of cells undergoing inactivation at that locus being highly gene-specific. We wished to determine if all cells in a population that silence one locus for a given gene, all silence the same locus; that is, is there a preference for which allele is inactivated. To this end, 7 highly expressed X-linked genes *(ACO1, APC, CRIM1, MLLT3, MPDZ, NFIB* and *RFX3*) were selected for which single nucleotide polymorphisms (SNPs) could be used to identify the expression of two different alleles. Using the FISH data of Livernois et al., (2013), that determined for a given gene the percentage of cells within a population that express one or both alleles, we were able to calculate the expected percentage of transcripts that would be from alleles ‘A’ and ‘B’ if all of the cells expressing just one allele (as identified by Livernois et al., 2013) expressed the same allele. Using our RNAseq data we calculated the percentage of transcripts representing alleles ‘A’ and ‘B’ for each gene. Observed percentages were compared to expected percentages and two-tailed P values were calculated using a chi squared test with 1 degree of freedom.

### Luciferase Reporter Assays

Luciferase reporter assays were used to confirm the absence of a functional ESRRB-SOX2 motif in the promoter of platypus *DAX1.* Constructs containing the platypus, mouse or ‘corrected’ platypus (where the murine Sox2 binding site was inserted into the platypus *DAX1* promoter adjacent to the ESRRB binding site with an intervening 5 nucleotides, as is the case in the mouse *Dax1* promoter) *DAX1* promotor were generated by Life Technologies Australia Pty Ltd (Fig. S1). Each construct contained 3 repeats of the ESRRB-SOX2 binding motif (Fig. S1). Sequences were subcloned into the pGL4.23[luc2/minP] vector (Promega) and each construct was transfected into mouse ESCs using Fugene-6 (Roche). Luciferase reporter assays were performed using the Luciferase Assay System (Promega) according to the manufacturer’s instructions. The β-Galactosidase Enzyme Assay System (Promega) was used as a control for transfection efficiency between wells, following the manufacturer’s protocol. Two preparations of each construct (‘a’ and ‘b’) were transfected; each transfection was repeated four times and duplicate measurements were made for each transfection. The data is presented as mean luciferase activity ± standard deviation.

## RESULTS

### Generation of Platypus iPSCs

Platypus fibroblast cultures were established from a punch biopsy of the toe webbing of an adult female platypus. Fibroblast cultures at passage 6 were reprogrammed into piPSCs using lentiviral delivery of human *OCT4, SOX2, KLF4, cMYC, NANOG* and *LIN28*. Reprogramming of platypus fibroblasts with human reprogramming factors was found to be exceptionally inefficient when compared to that of other mammals. PiPSCs form domed colonies consisting of cells that have a typical pluripotent stem cell morphology with a high nuclear to cytoplasmic ratio (Fig. 1*A*) and are karyotypically normal (Fig. 1*B*).

**Figure 1.**
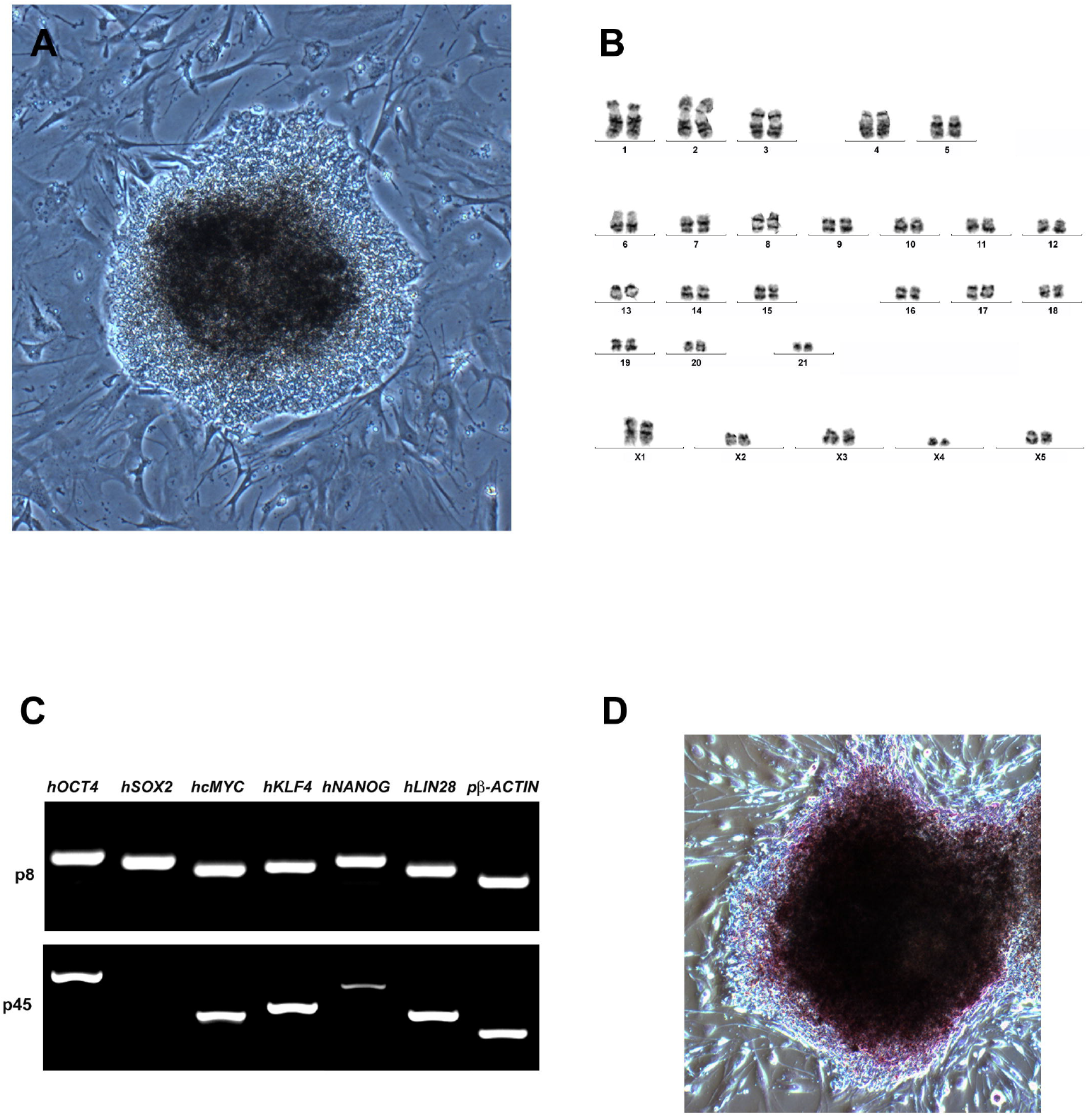
Platypus iPSCs have a characteristic pluripotent stem cell morphology *(A),* a large nuclear to cytoplasmic ratio, and form domed colonies. *(B)* piPSCs are karyotypically normal with 21 pairs of autosomes and 5 pairs of X chromosomes as is seen in female platypus. *(C)* At passage 8 all 6 of the human transgenes were still expressed in the piPSCs; at passage 45 *hSOX2* was silenced and expression of *hNANOG* was significantly reduced, while the remaining transgenes were still expressed. *(D)* piPSCs have robust alkaline phosphatase activity.

At passage 8, all six transgenes *(hOCT4, hSOX2, hcMYC, hKLF4, hNANOG* and *hLIN28)* were robustly expressed (Fig. 1*C*). By passage 45, silencing of *hSOX2* and a significant decrease in the expression of *hNANOG* were observed; however, expression of *hOCT4, hcMYC, hKLF4* and *hLIN28* persisted.

Platypus iPSCs possess alkaline phosphatase activity (Fig. 1*D*) and immunocytochemistry confirmed that they produce the pluripotency-associated cell surface markers SSEA1, SSEA4, TRA1-60 (Fig. 2A, *B* and C).

**Figure 2.**
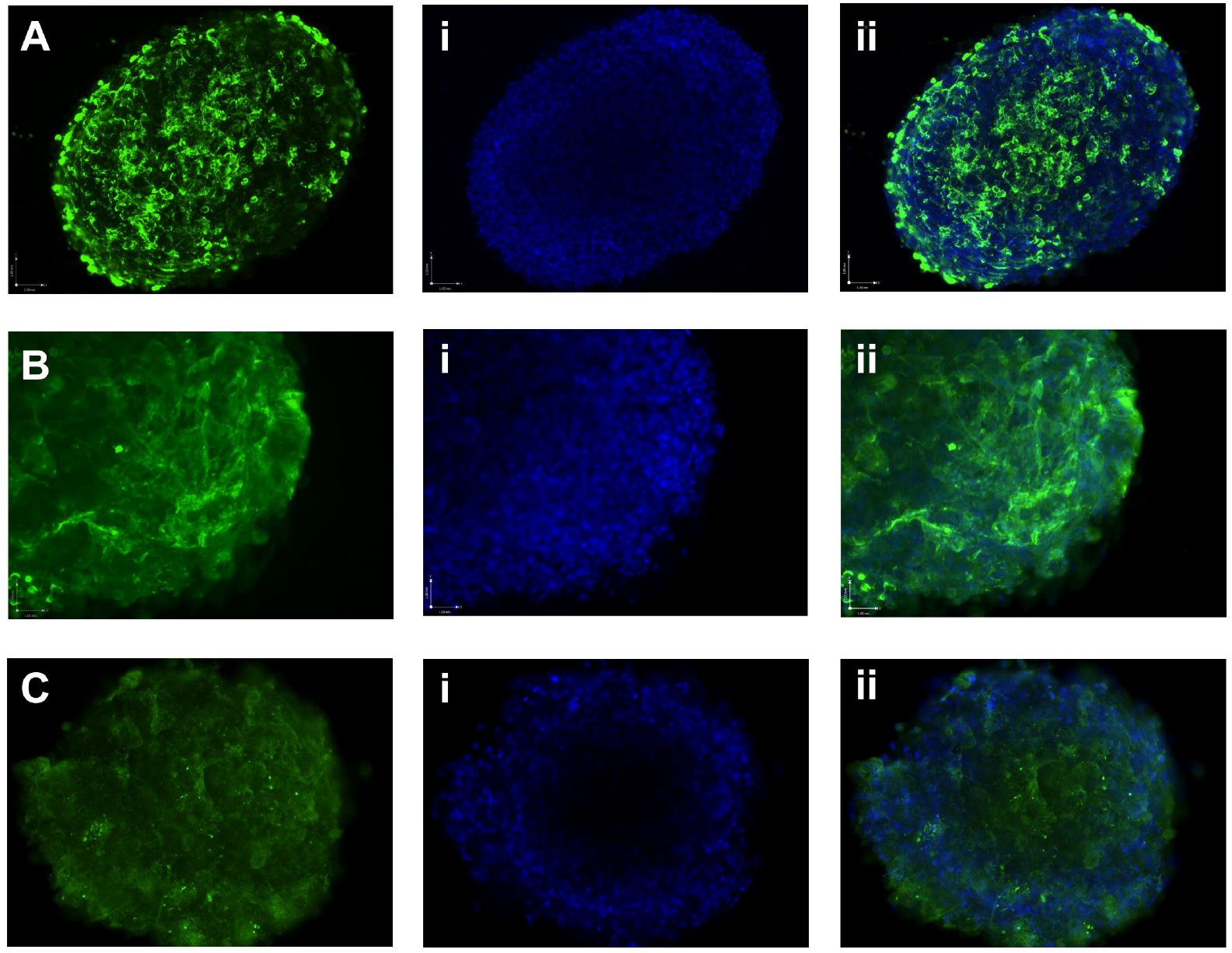
Platypus iPSCs express pluripotency-associated cell surface markers. *(A)* SSEA1. *(B)* SSEA4. *(C)* TRA1-60.

### Differentiation in Embryoid Bodies and In Vitro Teratomas

To assess the differentiation potential of the piPSCs we employed the embryoid body (EB) differentiation assay. PiPSCs readily form EBs in vitro (Fig. 3*A*) that after 6 weeks express the mesoderm marker *PAX2,* endoderm marker *GATA6* and the ectoderm marker *βIII-TUBULIN* (Fig. 3*B*) providing evidence of pluripotency of the piPSCs.

**Figure 3.**
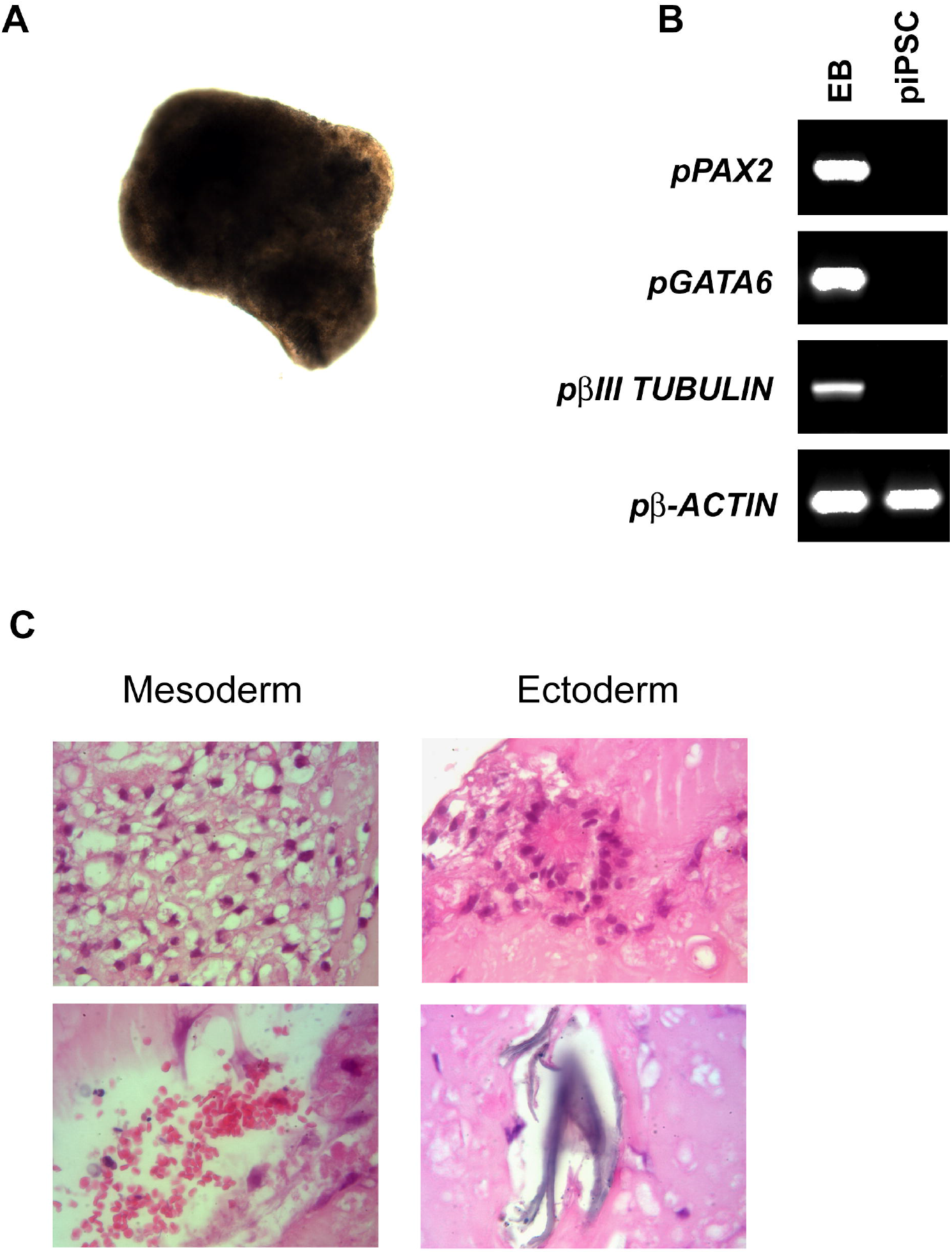
Platypus iPSCs form all three germ layers in vitro. *(A)* piPSCs readily form embryoid bodies (EBs) in vitro. *(B)* After 6 weeks in culture, EBs express the mesoderm marker *PAX2,* endoderm marker *GATA6* and the ectoderm marker *βIII-TUBULIN. (C)* After 8 weeks in culture, piPSC-derived in vitro teratomas contain adipose tissue and erythrocytes from mesoderm, and neural rosettes and keratin from ectoderm.

The most stringent test of pluripotency is the teratoma assay in immunodeficient mice. However, platypus iPSCs are maintained at 32°C, the core body temperature of the platypus, and do not survive at the mouse body temperature of 37°C. Consequently, this precluded the use of mice for the generation of conventional teratomas. We therefore used a recently developed in vitro differentiation assay that allows for the generation of mature tissues at 32°C (Whitworth et al. 2012; Whitworth et al. 2014a; Whitworth et al. 2014b; Weeratunga et al. 2018). PiPSC-derived in vitro teratomas contain adipose tissue and erythrocytes from mesoderm (Fig. 3*C*), and neural rosettes and keratin from ectoderm (Fig. 3*C*). No endoderm derivatives were identified. We have previously used this methodology with iPSCs from dog (Whitworth et al. 2012; Whitworth et al. 2014a), horse (Whitworth et al. 2014b) and Tasmanian devil (Weeratunga et al. 2018), and while in all of these studies endoderm derivatives were present, they occurred at a significantly lower frequency than derivatives from mesoderm and ectoderm. Thus, the lack of endoderm-derived tissues in platypus in vitro teratomas likely reflects a limitation of this assay rather than the piPSCs themselves, especially since *GATA6* expression is detected in EBs.

### Platypus iPSCs are LIF-dependent While bFGF Stimulates Their Proliferation

Since little is known about early platypus embryos, and stem cells from this species have not been established, we next wished to determine whether LIF and/or bFGF, two molecules that are key for maintaining pluripotency in mouse and human pluripotent stem cells, respectively, are required for the maintenance of pluripotency and/or proliferation in the piPSCs. To this end we cultured the piPSCs on mouse embryonic fibroblast (MEF) feeder layers in medium containing either LIF or bFGF, or both LIF and bFGF. Cells cultured on MEFs with bFGF alone, in the absence of LIF, failed to proliferate and eventually died, indicating LIF-dependency for the maintenance of pluripotency. Cells cultured with LIF alone did not proliferate as rapidly as cells cultured with both LIF and bFGF. To quantify this effect we performed immunocytochemistry using an antibody to phospho-histone H3 (Ser10) which identifies cells undergoing mitosis. PiPSCs cultured with both LIF and bFGF had a two-fold higher number of mitotic figures per HPF (12.93 ± 0.78 SEM) (Fig. 4*A* and *B*) than those cultured with LIF alone (6.65 ± 0.64 SEM, P < 0.0001) (Fig. 4*A* and *B*), indicating a role for bFGF signaling in the proliferation of piPSCs.

**Figure 4.**
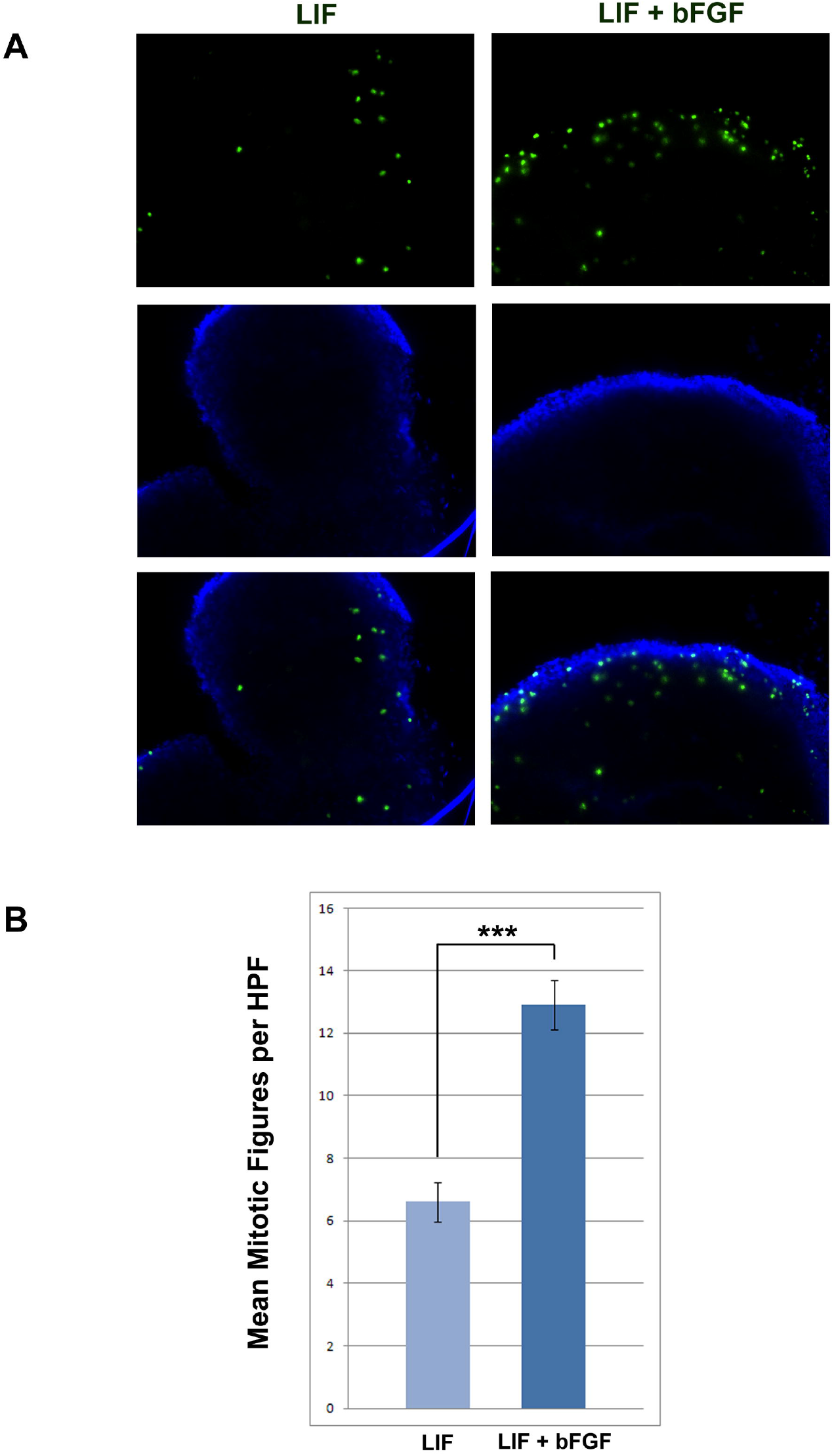
Platypus iPSCs are LIF-dependent but bFGF signaling stimulates proliferation. *(A)* piPSCs cultured with LIF alone have fewer mitotic figures (green immunofluorescence) than those cultured with both LIF and bFGF. *(B)* Graphic representation of the mean number of mitotic figures per high power field (HPF) ± s. e. m. for piPSCs maintained in medium with LIF or LIF and bFGF. piPSCs cultured with LIF and bFGF have a greater number of mitotic figures per HPF than those cultured with LIF alone. ***, P < 0.0001.

### Deep Sequencing of the piPSC Transcriptome Reveals a Unique Monotreme Pluripotency Signature

RNA sequencing data of the piPSC and platypus fibroblast transcriptomes revealed upregulated expression (FC_log_2_ ≥ 2) of many of the eutherian pluripotency factors in piPSCs, as compared to fibroblasts, including the core pluripotency factors *OCT4, SOX2* and *NANOG,* in addition to *DNMT3A, DNMT3B, FOXO1, GBX2, JARID2, LEF1, LEFTY1, LIF, LIFR, LIN28B, N-MYC, NOTCH1, NR6A1, SALL1, SALL4, SF1, SPP1, TCFAP2C* and *ZIC5* (Fig. 5*A* and for the full list see Dataset S1). Remarkably, *DAX1/NR0B1, ESRRB, NR5A2, PRDM14* and *TBX3,* all of which are known to play key roles in establishing and/or maintaining pluripotency in eutherians, are not expressed in platypus iPSCs (FPKM < 1) (Fig. 5*A* and Dataset S1).

**Figure 5.**
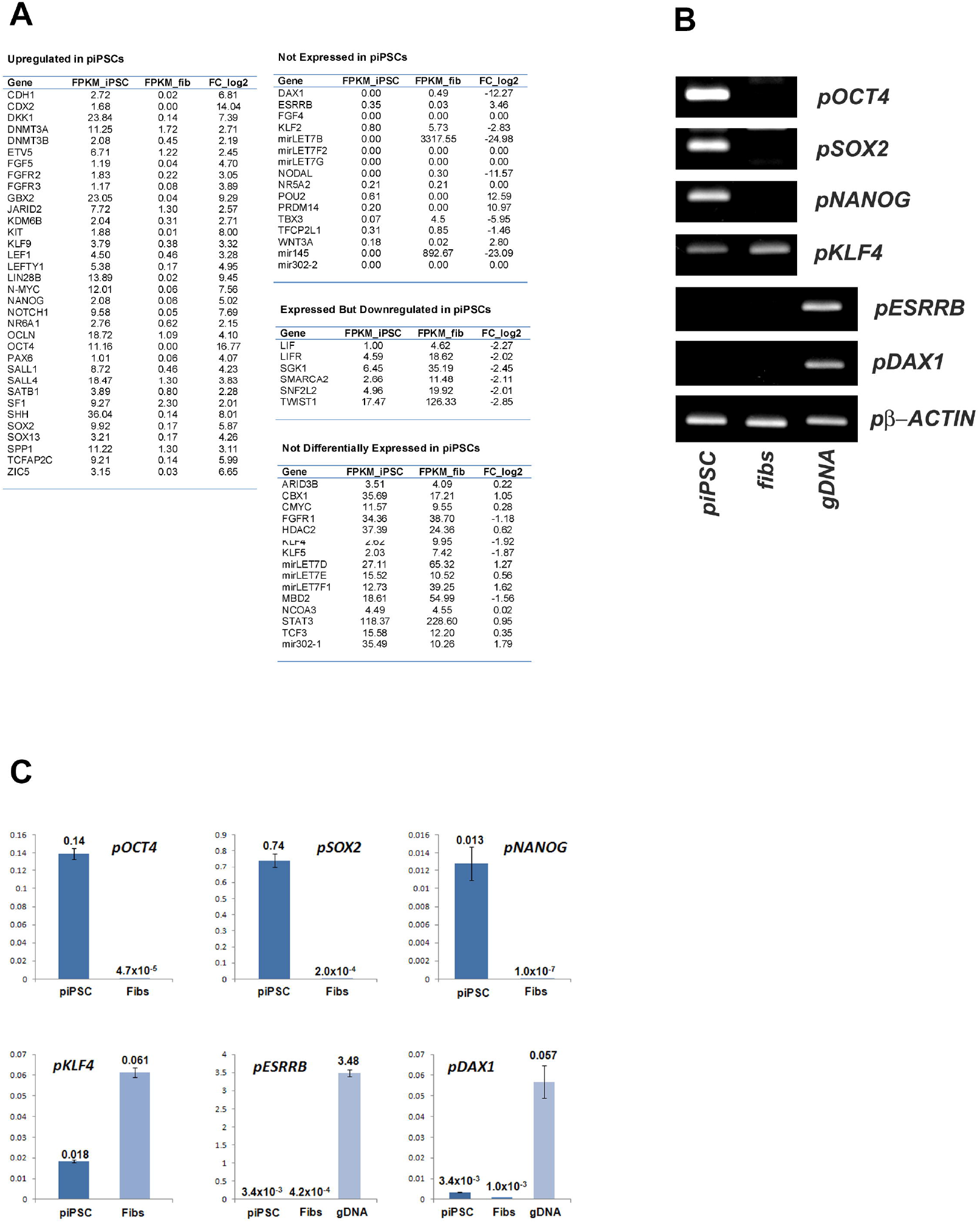
Deep sequencing of the piPSC transcriptome identified a unique monotreme pluripotency signature. *(A)* RNA sequencing data of the piPSC and platypus fibroblast transcriptomes revealed upregulated expression (FC_log_2_ > 2) in piPSCs, as compared to fibroblasts, of many of the pluripotency factors identified in eutherians including the core pluripotency factors *OCT4, SOX2* and *NANOG.* Remarkably, *DAX1/NR0B1, ESRRB, NR5A2, PRDM14* and *TBX3* are not expressed in platypus iPSCs (FPKM < 1). *(B)* Using RT-PCR we confirmed that platypus *OCT4, SOX2* and *NANOG* are expressed in piPSCs and not in fibroblasts. In agreement with the RNA sequencing data, *pKLF4* is expressed by both piPSCs and fibroblasts. Neither *pESRRB* nor *pDAX1* expression could be detected in either piPSCs or fibroblasts. *(C)* Quantitative RT-PCR confirmed expression of *pOCT4, pSOX2* and *pNANOG* in piPSCs, but not fibroblasts, and the expression of *pKLF4* in both piPSCs and fibroblasts. Expression levels of both *pESRRB* and *pDAX1* are at background levels confirming the very low numbers of transcripts identified by RNA sequencing.

Six members of the *LET7* family of microRNAs (miRNAs) have been identified in the platypus genome: *LET7B, LET7D, LET7E, LET7F1, LET7F2* and *LET7G.* Of these, *LET7F2* and *LET7G* are not expressed in piPSCs or fibroblasts, while *LET7B* is found at very high levels in fibroblasts (FPKM = 3317.55) but is not detected in piPSCs (FPKM = 0.00) (Fig. 5*A* and Dataset S1). Other *LET7* miRNAs, *LET7D, LET7E* and *LET7F1,* are not differentially expressed between piPSCs and fibroblasts (Fig. 5*A* and Dataset S1). *miR-145,* a miRNA associated with the transition out of pluripotency in human pluripotent stem cells (Xu et al. 2009), is transcribed at very high levels in adult fibroblasts (FPKM = 892.67) but is not expressed in piPSCs (FPKM = 0.00) (Fig. 5*A* and Dataset S1). These data therefore suggest that *LET7*-mediated regulation of puripotency is a regulatory layer most likely acquired during eutherian evolution.

In agreement with the observed LIF dependency, piPSCs express both *LIF* and *LIF receptor (LIFR)* (FPKM = 1 and 4.59, respectively). Surprisingly, both are also expressed in the platypus fibroblasts (Fig. 5*A* and Dataset S1). In support of their proliferative response to bFGF, piPSCs express the FGF receptors *FGFR1, FGFR2* and *FGFR3* (FPKM = 34.36, 1.83 and 1.17, respectively) (Fig. 5*A* and Dataset S1).

Several genes associated with the eutherian plurinet were found to be expressed at similar levels (FC_log_2_ > -2 or < 2) in both piPSCs and adult fibroblasts including *c-MYC, KLF4, KLF5, NCOA3, STAT3, TCF3* and the microRNA *miR302-1* (Fig. 5*A* and Dataset S1).

Given the surprising finding that neither *DAX1/NR0B1* nor *ESRRB* are expressed in platypus iPSCs we sought to confirm these data using qRT-PCR. The expression of platypus *OCT4, SOX2* and *NANOG* is clearly detected in piPSCs and not in fibroblasts (Fig. 5*B* and *C*). In contrast, but in agreement with the RNA sequencing data, *KLF4* is expressed by both piPSCs and fibroblasts (Fig. 5*B* and *C*). Neither *ESRRB* nor *DAX1* expression could be detected by qRT-PCR beyond background levels in either piPSCs or fibroblasts (Fig. 5*B* and *C*), confirming the very low numbers of transcripts identified by RNA sequencing.

Prompted by these novel findings in the platypus plurinet we next compared the pluripotency networks of chicken (as determined from chicken ESCs; Jean et al., 2015), mouse (iPSCs; Liu et al., 2014) and pig (iPSCs; Liu et al., 2014) with that of the platypus. *NANOG* and *OCT4,* in addition to *DNMT3B, TFAP2C, NR6A1* and *OCLN,* are amongst the 39 genes shared by all four species (Fig. 6 and Dataset S2), supporting their position as ancestral pluripotency genes likely shared by the common ancestor of birds and mammals.

**Figure 6.**
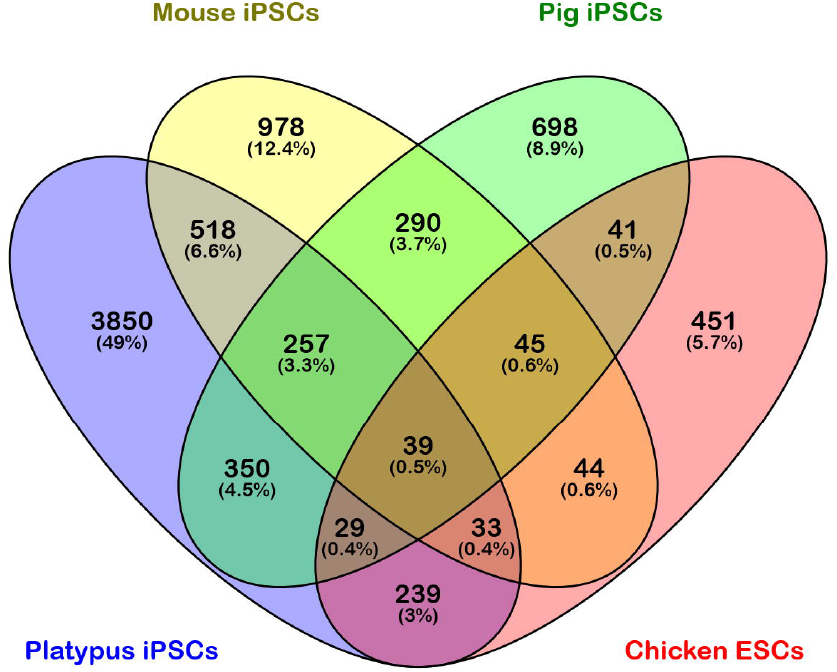
*NANOG* and *OCT4* are amongst the 39 pluripotency-associated genes that are expressed by platypus and chicken, in addition to mouse and pig, and so represent ancestral pluripotency factors likely shared by the common ancestor of birds and mammals. Platypus and chicken share a further 239 unique genes which may include genes that are required for pluripotency in birds and monotremes, but which have lost this role in therian mammals.

Platypus and chicken share a further 239 unique genes (Fig. 6 and Dataset S2) which may include genes that are required for pluripotency in birds and monotremes but which have lost this role in therian mammals. Platypus, mouse and pig have 257 genes in common, including *APOE, BRCA1, DNMT1, JARID2, POU2F1, PTCH1* and *SOX13* (Fig. 6 and Dataset S2), which are likely ancestral members of the mammalian plurinet.

### *DAX1* Acquired SOX2 Responsiveness During Therian Evolution

DAX1 is expressed in eutherian pluripotent stem cells where it has been shown to be essential for maintaining pluripotency (Khalfallah et al., 2009; Hutchins et al., 2013). Thus, in an attempt to understand the lack of *DAX1* expression in piPSCs we examined the promoter region which, in eutherians, has been shown to contain a composite ESRRB-SOX2 DNA binding site that is essential for transcription of *Dax1* in mouse ESCs (Hutchins et al., 2013). Significantly, activity of the dual binding motif is lost if either the Esrrb or Sox2 binding site is disrupted (Hutchins et al., 2013). While the ESRRB DNA binding site is conserved amongst all 3 mammalian subclasses - monotremes (platypus), marsupials (opossum and wallaby) and eutherians (cow, pig, mouse, dog and human) - the SOX2 binding site is absent in the platypus (Fig. 7). Curiously, in both marsupial species examined, there are two possible SOX2 binding sites that are conserved, but these are 17 and 21 base pairs downstream from the ESRRB binding site, making it unlikely that ESRRB and SOX2 co-assemble to form an ESRRB-SOX2-DNA ternary complex (Fig. 7).

**Figure 7.**
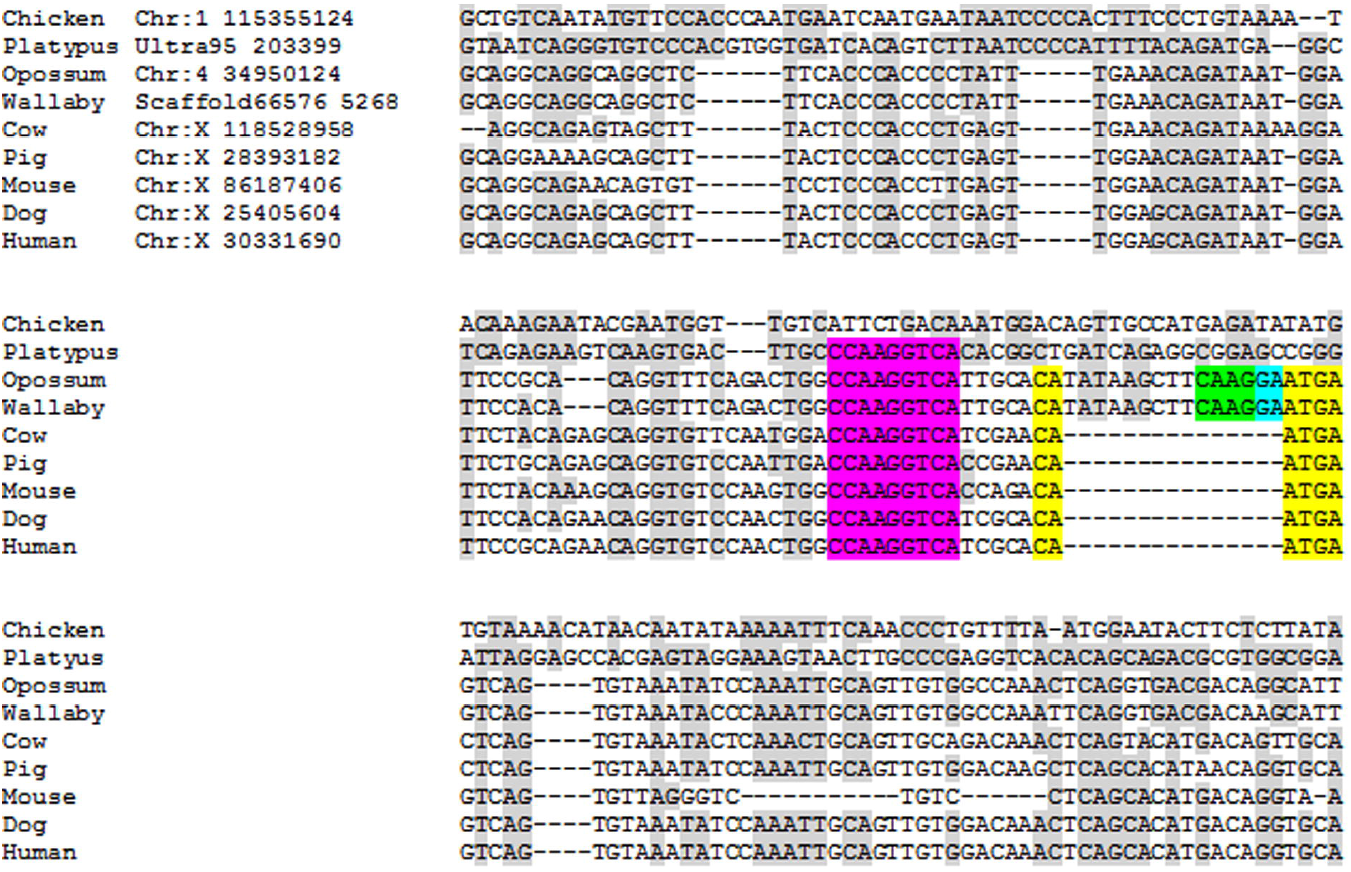
*DAX1* acquired SOX2 responsiveness during therian evolution. While the ESRRB DNA binding site (magenta) is conserved amongst all 3 mammalian subclasses - monotremes (platypus), marsupials (opossum and wallaby) and eutherians (cow, pig, mouse, dog and human) - the SOX2 binding site (yellow) is absent in the platypus. In both marsupial species examined, there are two possible SOX2 binding sites that are conserved (green and blue), that are 17 and 21 base pairs, respectively, downstream from the ESRRB binding site, making it unlikely that ESRRB and SOX2 co-assemble to form an ESRRB-SOX2-DNA ternary complex. It is possible that the eutherian SOX2 DNA binding site (yellow) has been created via an excisional event, removing approximately 15 base pairs of intervening sequence that is present in the marsupial genome.

### Platypus lacks a functional ESRRB-SOX2 co-binding motif in its *DAX1* promoter

When transfected into mouse ESCs, the platypus *DAX1* promotor construct produced only background levels of luciferase activity (a: 171.73 ± 19.16; b: 168.93 ± 5.24) (Fig. 8). In contrast, the mouse *Dax1* promotor construct displayed high levels of luciferase activity (a: 8124.65 ± 171.53; b: 7497.84 ± 162.42) (Fig. 8). Our attempt at ‘correcting’ the platypus *DAX1* promotor by inserting the murine Sox2 binding site 5 bp downstream of the ESRRB binding motif did not result in increased luciferase activity as compared to the platypus *DAX1* promotor construct (a: 244.81 ± 34.79; b: 134.99 ± 30.67) (Fig. 8).

**Figure 8.**
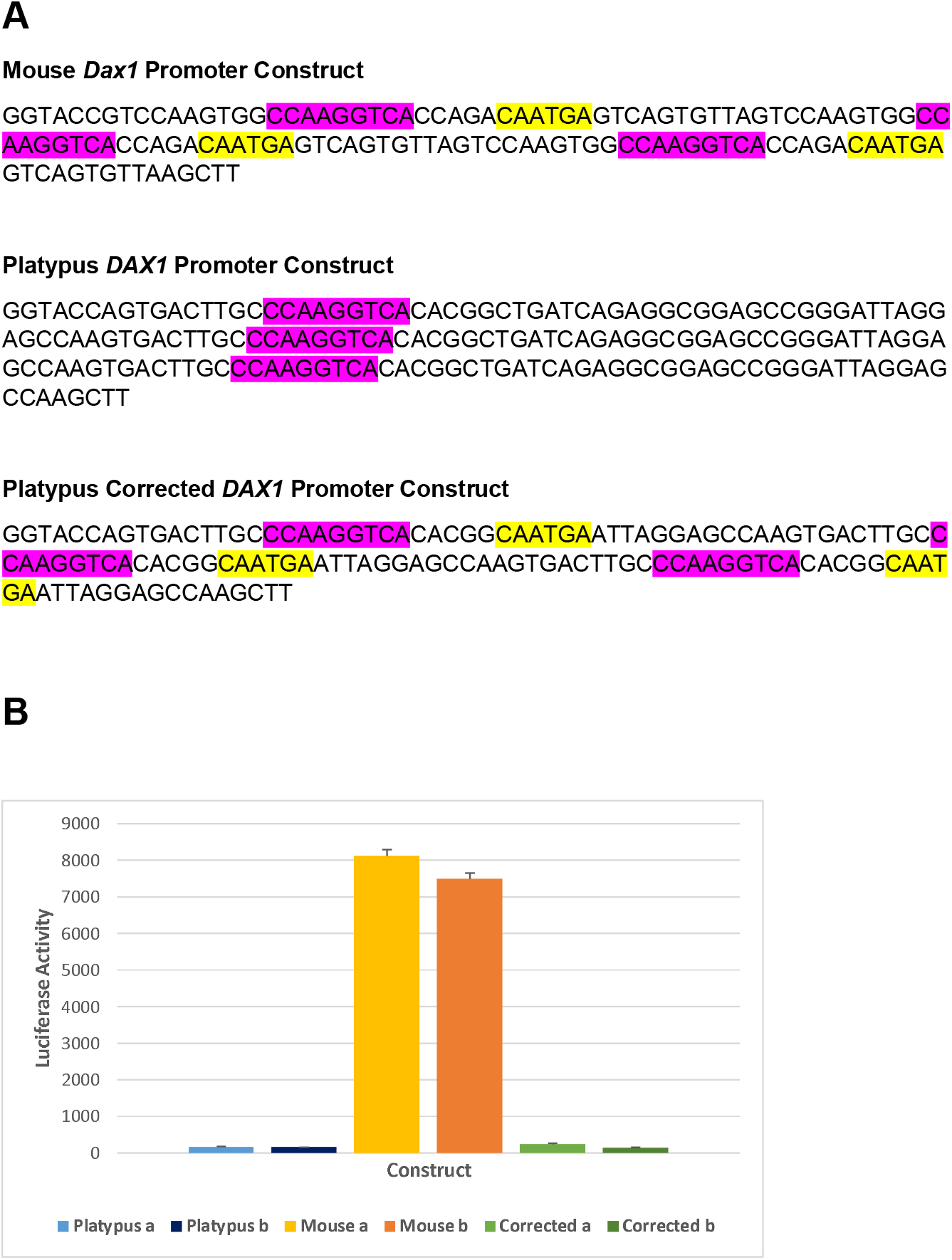
*(A)* Sequences of constructs containing the platypus, mouse or ‘corrected’ platypus *DAX1* promotor sequence. The ESRRB and SOX2 binding motifs are shown in pink and yellow, respectively. *(B)* When transfected into mouse ESCs, the platypus *DAX1* promotor construct produced only background levels of luciferase activity while the mouse *Dax1* promotor construct displayed high levels of luciferase. The ‘corrected’ platypus *DAX1* promotor that contained the murine Sox2 binding site did not result in increased luciferase activity as compared to the platypus *DAX1* promotor construct.

### Expression Ratios of X Chromosomes to Autosomes Indicate a Lack of Global Upregulation of X-Linked Genes

The median expression level of X-linked genes in the fibroblasts was 7.579, and that of autosomal genes was 7.000, giving an expression ratio of X chromosomes to autosomes (X_1-5_ X_1-5_:AA) of 1.08 which does not support upregulation of X-linked genes. Similarly, the median expression level of X-linked genes in the piPSCs was 5.703, as compared to a median expression level of 6.637 for the autosomal genes, giving an X_1-5_ X_1-5_:AA of 0.85 which also does not suggest an upregulation of X-linked genes.

### Platypus randomly inactivates both maternal and paternal loci within a population of cells

Using single nucleotide polymorphisms (SNPs) to distinguish between two alleles for each of the 7 X-linked genes examined, and comparing the actual ratio of transcripts A to B to the expected ratio, as predicted by the FISH data of Livernois et al., (2013), we are able to demonstrate that for 6 of the 7 genes both alleles are expressed at close to a 50:50 ratio (Table 1). The exception is NFIB where 61% of transcripts correspond3d to allele A and 39% to allele B (Table 1). However, for all 7 genes, the expected and observed results were statistically significantly different indicating that when cells inactivate an X-linked gene, there is no consistency between cells as to which locus is inactivated; thus, the platypus does not appear to preferentially inactivate either the maternal or paternal X-linked gene.

**Table 1:** Platypus randomly inactivates both maternal and paternal loci within a population of cells. * FISH data from Livernois et al., 2013. # Where allele A is the transcript detected in cells with 2 signals and in cells with 1 signal; allele B is the transcript also detected in cells with 2 signals.

| Gene | % Nuclei with 2 signals * | % Nuclei with 1 signal * | Expected % of transcripts allele A # | Expected % of transcripts allele B # | Observed % of transcripts allele A | Observed % of transcripts allele B | P-value |
| --- | --- | --- | --- | --- | --- | --- | --- |
| *ACO1* | 57 | 39 | 63 | 37 | 50 | 50 | 0.001 |
| *APC* | 50 | 50 | 67 | 33 | 52 | 48 | 0.001 |
| *CRIM1* | 20 | 80 | 83 | 17 | 53 | 47 | 0.0001 |
| *MLLT3* | 38 | 58 | 72 | 28 | 58 | 42 | 0.001 |
| *MPDZ* | 59 | 38 | 62 | 38 | 50 | 50 | 0.01 |
| *NF1B* | 31 | 62 | 78 | 22 | 61 | 39 | 0.0001 |
| *RFX3* | 45 | 51 | 68 | 32 | 53 | 47 | 0.001 |

## DISCUSSION

In this paper we report the generation of the first pluripotent stem cells from a monotreme. Platypus iPSCs, like the platypus itself and its genome, appear to reflect both derived and ancestral characteristics. The platypus plurinet consists of many of the genes known to play important roles in eutherian pluripotency, including the core pluripotency factors *OCT4, SOX2* and *NANOG.*

*OCT4,* also known as *POU5F1,* is a class V POU family transcription factor that, along with *SOX2* and *NANOG,* is a core regulator of pluripotency in eutherians. This study has confirmed the ancestral role of *OCT4* in mammalian pluripotency as piPSCs, but not fibroblasts, express *OCT4.* The chicken lacks an *OCT4* orthologue, but the paralogue to *OCT4, POU2 (POUV),* is expressed in chicken ESCs (Lavial et al., 2007; Jean et al., 2015). *OCT4* arose by duplication of *POU2* early in the evolution of tetrapods, and while *POU2* has been lost from the genome of eutherian mammals, it persists in the genomes of marsupials and the platypus (Niwa et al. 2008; Frankenberg et al. 2010). In piPSCs, there is some transcription of *POU2* but at levels below the threshold considered to represent expression. We have recently generated iPSCs from a marsupial, the Tasmanian devil, where we observed robust expression of both *POU2* and *OCT4* (Weeratunga et al., 2018). During marsupial development, *OCT4* is expressed by cells of the epiblast (the equivalent of the eutherian inner cell mass), hypoblast and trophectoderm of pre-implantation embryos, while expression of *POU2* is restricted to the epiblast (Woranop et al., 2015). Given that both marsupials and monotremes possess the *POU2* gene, and that it appears to play a central role in marsupial pluripotency, it is surprising that *POU2* is not expressed in the piPSCs. An explanation for this may be that the human *OCT4* transgene was still being expressed in the piPSCs, which may have a down regulatory effect on *POU2* expression.

Several genes which are known to be involved in eutherian pluripotency are expressed at comparable levels in both piPSCs and adult fibroblasts. One such gene is *KLF4,* which is a key regulator of pluripotency in eutherians and is one of the factors commonly used for the reprogramming of cells into iPSCs across a range of eutherian species (Takahashi and Yamanaka 2006; Takahashi et al. 2007; Whitworth et al. 2012; Whitworth et al. 2014b; Weeratunga et al., 2018). While *KLF4* is present at significant levels in piPSCs, as would be expected of a regulator of pluripotency, its levels are comparable to those detected in the adult fibroblasts. *KLF4* is highly expressed in the epidermis of eutherians (Garrett-Sinha et al. 1996; Segre et al. 1999) and has also been detected, albeit at lower levels, in human dermal fibroblasts maintained in culture (Aasen et al. 2008), thus possibly accounting for the significant levels of *KLF4* expression that we see in platypus dermal fibroblasts.

*C-MYC* is similarly expressed at comparable levels in both fibroblasts and piPSCs and, as with *KLF4,* is also expressed within the epidermis of adult skin (Hurlin et al. 1995; Bull et al. 2001) and cultured dermal fibroblasts (Aasen et al. 2008). In contrast, the *c-MYC* paralogue, *N-MYC,* is robustly expressed in piPSC but is not expressed in fibroblasts. Both *c-Myc* and *N-Myc* have been shown to be essential for maintaining pluripotency and stimulating proliferation in mouse iPSCs (Smith et al. 2010). It is likely that *c-Myc* exerts its control over cell proliferation via transactivation of *Lin28b* (Chang et al. 2009) which, along with its paralogue *Lin28a,* regulates the self-renewal of pluripotent stem cells by suppressing the *let7* family of miRNAs which are thought to promote differentiation in pluripotent stem cells (Chang et al. 2009; Melton et al. 2010; Suh et al. 2010). A feedback loop exists between *Lin28a* and *Lin28b* since *let7* miRNAs can, in turn, repress the expression of *Lin28a* and *Lin28b* (Shyh-Chang and Daley 2013). In the platypus, we see high levels of *LIN28B* expression in the piPSCs, but not in fibroblasts. Unfortunately, the platypus *LIN28A* gene is not annotated.

Six members of the *LET7* family have been identified in platypus: *LET7B, LET7D, LET7E, LET7F1, LET7F2* and *LET7G.* Of these, *LET7F2* and *LET7G* are not expressed in piPSCs or fibroblasts, while *LET7B* is found at very high levels in fibroblasts but is not detected in piPSCs. *LET7D, LET7E* and *LET7F1* are expressed at similar levels in piPSCs and fibroblasts. These disparate expression profiles of the various members of the *LET7* family are at odds with what has been demonstrated in the mouse where none of the *let7* miRNAs are present in mESCs (Thomson et al. 2004). However, it is important to note that miRNAs are transcribed as primary miRNAs (pri-miRNAs) which are subsequently processed in the nucleus to form pre-miRNAs which, in turn, give rise to the mature miRNA (Roush and Slack 2008). Although mature *let7* miRNAs have not been detected in mouse ESCs, pri-let7 miRNAs are transcribed (Thomson et al. 2004) and pre-let7 transcripts are present in the cytoplasm (Ryback et al. 2008). Thus, the transcripts that we are detecting in the platypus may constitute pri-, pre- or mature *LET7* miRNAs, or a combination of all 3, and so it is not possible to determine if the *LET7* miRNAs detected in the platypus piPSCs are the functional mature form. It is perhaps also significant that while *LET7* miRNAs are known to downregulate expression of *LIN28A* and *LIN28B,* significant levels of *LIN28B* expression are present in piPSCs despite the presence of *LET7* transcription (Roush and Slack 2008).

An additional miRNA known to play an important role in facilitating the shift out of pluripotency is *miR-145,* which has been shown to downregulate the expression of *OCT4, SOX2* and *KLF4,* and which is itself downregulated by OCT4 (Xu et al. 2009). The platypus *miR-145* is transcribed at high levels in adult fibroblasts but is not expressed in piPSCs, suggesting a similar role for this miRNA in platypus pluripotent stem cells as in those of eutherians.

Leukaemia inhibitory factor (LIF) is sufficient to maintain pluripotency and self-renewal in mouse ESCs and iPSCs (Smith et al. 1988; Williams et al. 1988), and appears to be an ancestral requirement of pluripotent cells since LIF signaling is also necessary for pluripotency and self-renewal in avian ESCs (Pain et al. 1996; Lavial et al. 2007). In the mouse, LIF signals via the activation of *Stat3* (Niwa et al. 1998). Platypus iPSCs are LIF-dependent and express both *LIF* and the *LIF receptor (LIFR),* in addition to *STAT3.* Recently, studies in the mouse have identified *Gbx2* as a downstream target for LIF/STAT signaling in mESCs (Tai and Ying 2013). Over-expression of *Gbx2* is sufficient to maintain mouse *Stat3* null ESCs and has also been shown to be a marker specific for naïve ESCs as compared to epiblast stem cells which exist in a more primed state of pluripotency (Tai and Ying 2013). Significant levels of *GBX2* expression are seen in piPSCs, but not in fibroblasts, indicating that the LIF/STAT3/GBX2 signaling pathway is an ancestral component of the pluripotency network.

While piPSCs are LIF-dependent, the addition of exogenous bFGF (FGF2) to cells in culture induces a proliferative effect and, in keeping with this observed effect, the FGF receptor *FGFR1* is also expressed in piPSCs, as is seen in human ESCs (Coutu and Galipeau 2011). Other components of the FGF/FGFR signaling pathway are similarly expressed in piPSCs including *FRS2, GRB2, GAB1* and *MAPK1* (Lanner and Rossant 2010). Unfortunately, the gene for FGF2 is not annotated in the platypus and so we were unable to examine endogenous levels of *FGF2* expression. While FGF2 is the predominant FGF expressed in human pluripotent cells, mouse ESCs express high levels of FGF4 which likely also signals through FGFR1 (Lanner and Rossant 2010). Platypus iPSCs do not express *FGF4.* The 3’ UTR of *FGF4* in eutherian mammals, the platypus, and the chicken, contains a SOX2-OCT4 binding motif (Fernandez-Tresguerres et al. 2010) that has been shown in the mouse to drive high levels of *Fgf4* transcription (Remenyi et al. 2003). While the SOX2 binding motif in the enhancer is identical between platypus and eutherian mammals, the OCT4 binding site shows a nucleotide substitution and a deletion of one base which is thought to disrupt OCT4 binding (Fernandez-Tresguerres et al. 2010). The chicken OCT4 binding motif is similarly disrupted and in an enhancer activity assay in mouse ESCs, the chicken enhancer was found to be ineffective at driving expression of the reporter (Fernandez-Tresguerres et al. 2010). *FGF4* is not expressed in chicken pluripotent stem cells (Fernandez-Tresguerres et al. 2010) which is consistent with the inability of the SOX2-OCT4 binding motif to drive *FGF4* expression in mESCs. Thus, it is possible that the lack of *FGF4* expression in piPSCs is similarly due to the proposed inactivity of the SOX2-OCT4 binding motif in the platypus *FGF4* enhancer.

*Esrrb* is an orphan nuclear receptor that is a downstream target of *Nanog* (Festuccia et al. 2012) and is both necessary, and sufficient, for self-renewal in mouse ESCs (Ivanova et al. 2006; Loh et al. 2006; Festuccia et al. 2012; Martello et al. 2012; Percharde et al. 2012). Studies by Martello and colleagues (2012) suggest that Esrrb functions in parallel, and shares significant functional redundancy, with the LIF/Stat3 pathway. Further, while *Nanog* is considered a core factor of the plurinet, and over-expression of *Nanog* can compensate for loss of *Lif* signaling (Chambers et al. 2003), *Esrrb* can functionally rescue self-renewal in *Nanog* null and *Lif* null mESCs (Martello et al. 2012) further illustrating the high degree of redundancy between *Nanog*, *Esrrb* and *Lif*. *ESRRB* is not expressed in platypus or Tasmanian devil iPSCs (Weeratunga et al., 2018), and taken together with the significant functional redundancy between *Esrrb, Nanog* and *Lif* in the mouse, this supports the suggestion that *Esrrb* has been conscripted into the plurinet relatively late in mammalian evolution.

Another member of the eutherian pluripotency network that is not expressed in piPSCs is the orphan nuclear receptor *DAX1,* also known as *NR0B1.* In the mouse, *Dax1* loss-of-function in mESCs results in loss of pluripotency and subsequent differentiation (Clipsham et al. 2004; Niakan et al. 2006; Khalfallah et al. 2009). Hutchins and colleagues (2013) described the presence of an Esrrb-Sox2 binding motif in the promoter of murine *Dax1* which is specifically active in mESCs. Importantly, each of the Esrrb and Sox2 binding sites is independently necessary for *Dax1* expression since the binding of the transcription factors to their respective sites forms an Esrrb-Sox2-DNA ternary complex that mediates transcription (Hutchins et al. 2013). An alignment of this motif in the promoters of *DAX1* from several eutherian species, two species of marsupial and the platypus, reveals that while the ESRRB site is conserved, the SOX2 site is absent in the platypus. In both marsupial species examined, there are two possible SOX2 binding sites that are conserved, but these are 17 and 21 base pairs downstream from the ESRRB binding site, making it unlikely that ESRRB and SOX2 co-assemble to form an ESRRB-SOX2-DNA ternary complex as is the case in eutherians where only 5 base pairs separate the ESRRB and SOX2 binding sites. Furthermore, while marsupial iPSCs express *DAX1,* they do not express *ESRRB* (Weeratunga et al., 2018); thus, dual binding of ESRRB and SOX2 is not required for *DAX1* expression in marsupials, but binding of SOX2 alone may be sufficient. The absence of the SOX2 binding site in the platypus may therefore explain the absence of *DAX1* expression in piPSCs. Thus, the composite ESRRB-SOX2 binding site in the eutherian promoter of *DAX1* appears to have been created after the divergence of eutherians from the therian lineage via an excisional event that removed 15 base pairs of intervening sequence to bring the ESRRB and SOX2 binding sites within close proximity of each other.

Based on studies in the chicken, *POU2/OCT4* and *NANOG* appear to be the most ancestral of the core pluripotency factors, since both are expressed within chicken ESCs and the epiblast of pre-primitive streak-stage embryos (Lavial et al., 2007; Jean et al., 2015). The role of *SOX2,* however, is less clear. Expression of *SOX2* isn’t observed until the development of the neural plate at Hamburger-Hamilton stage 4 (Uchikawa et al. 2003; Fernandez-Tresguerres et al. 2010). In contrast, the closely related gene, *SOX3,* is co-expressed with *POU2/OCT4* and *NANOG* in pre-primitive-streak stage embryos (Jean et al., 2015). In vitro, *SOX2* expression has been detected within chicken ESCs, but is barely above background levels in blastodermal cells (cBCs), and is absent in primordial germ cells (PGCs), whereas *SOX3* is highly expressed within all three types of avian pluripotent stem cells (cESCs, cBCs and PGCs) (Jean et al., 2015). Hence, in the chicken, *SOX3* appears to play a more extensive role in pluripotency than *SOX2* (Fernandez-Tresguerres et al. 2010; Jean et al., 2015). Thus, within the 324 million years of independent evolution that separates birds and mammals from their nearest common ancestor, *SOX2* has replaced *SOX3* within the mammalian pluripotency pathway.

Ohno’s hypothesis of X chromosome dosage compensation proposes that there has been an upregulation in the expression of X-linked genes in both males and females in order to restore their expression to levels that existed before the differentiation of sex chromosomes, and to bring them back into parity with the expression levels of genes on the autosomes (Ohno, 1967). Using our RNA-seq data for the platypus fibroblasts and iPSCs we determined that for each cell type the ratio between the median transcription levels for all annotated X-linked genes across the five X chromosomes and those that are autosomal (X_1-5_X_1-5_:AA) is approximately 1.0, supporting the argument that there has been no global upregulation of X-linked genes in the platypus. Our data supports that of Julien et al. (2012) who also obtained a ratio of approximately 1.0 when comparing transcription of genes on X chromosome 5 to transcription from the autosomes.

Previous studies on the platypus have demonstrated that for any given X-linked gene, the proportion of nuclei within a population of cells that will have undergone silencing of one allele ranges from approximately 25-80%, with no gene displaying silencing of one allele in 100% of the cells examined (Deakin et al., 2008; Livernois et al., 2013). Thus, in monotremes, rather than inactivation of an entire X chromosome, there appears to be inactivation of X-linked genes on a gene-by-gene basis, with each gene expressed from one or both loci at a characteristic frequency (Livernois et al., 2013). This raises the question of whether for each gene, all cells that undergo silencing of one allele inactivate the same allele. We addressed this question using our RNA-seq data for the fibroblasts and found that for each of the X-linked genes examined, where 2 alleles could be identified using SNPs, the expected ratio of the two alleles if one allele was always preferentially inactivated was significantly different from the observed ratio which was, in most cases, approximately 50% for each allele. Livernois et al. (2013) found that in the case of neighbouring genes each cell inactivated loci on the one chromosome; however, their data did not enable them to determine if it was the same chromosome in each cell – that is, if the silencing is imprinted. Our data suggest that imprinting does not occur. While adjacent genes may be inactivated on the same chromosome, it is not always the same chromosome since even when 80% of cells are expressing only the one allele, approximately 50% of the transcripts are coming from each allele.

The pluripotency network of eutherian mammals has evolved through the co-option of genes and the development of new regulatory mechanisms. One key event in the evolution of mammalian pluripotency is the expanding role of SOX2 over SOX3, which occurred prior to 166 million years ago as we have shown in this study that SOX2 is indeed a constituent of the monotreme plurinet. Further support for the evolving role of SOX2 in mammalian pluripotency can also be found in the platypus where the absence of expression of *DAX1* and *ESRRB,* two key members of the eutherian plurinet, are likely due to an inability to respond to SOX2. A study by Buganim and colleagues (2012) in mouse iPSCs has placed *Sox2* at the start of a hierarchical cascade of gene expression that results in the acquisition of pluripotency. That is, the activation of *Sox2* appears to be the switch that activates the pluripotency circuitry. This stand-alone position of Sox2 within the plurinet is consistent with it having been recruited into the initial Oct4-Nanog machinery which persists in primordial germ cells. Pluripotency in the platypus reflects this evolving role of SOX2 in mammalian evolution and, in particular, the role of SOX2 in recruiting further members into the plurinet. Our data are consistent with the idea that repurposing of key transcription factors followed by secondary mutations in regulatory elements of other genes to create binding sites for such transcription factors may be a principle that has driven the evolution of gene regulatory networks.

---

**Supplementary Figure 1**. Constructs containing the platypus, mouse or ‘corrected’ platypus *DAX1* promotor sequence. Each construct contained 3 repeats (indicated by the green arrows) of the ESRRB (pink) and SOX2 (yellow) binding motifs.

**Where cells are expressing just one locus, it is not consistently the same locus but appears to be random with no preference for one allele over the other.**

